# A Condition-Informed Framework for Evaluating Temperature-Responsive Gene Sets Using Protein Interaction Data

**DOI:** 10.64898/2025.12.10.693541

**Authors:** Bradley Frishman, Jorge Gonzalez, Valery Forbes

**Affiliations:** College of Public Health and Health Professions, University of Florida, Gainesville, FL; Department of Mathematics, Embry-Riddle Aeronautical University, Daytona Beach, FL; Department of Biological Sciences, Florida Atlantic University, Boca Raton, FL

**Author notes:** Correspondence: *Bradley Frishman (*).

**Keywords:** Differentially Expressed Genes, Genomics, Housekeeping Genes, Protein-Protein Interactions, RNA-seq

## Abstract

Computational models of genomic responses to environmental perturbations identify genes underlying physiological adaptation, but their utility requires further evaluation. Often, gene activity reflects both adaptive mechanisms and noise. Here, we develop a framework that leverages public databases to support the subselection of biologically supported model genes involved in temperature responses. We test our framework on a model that identified differentially expressed genes (DEGs) and categorized some into Key-Response, Treatment-Specific, Noisy, and Support groups based on inter-individual gene expression variability before and after treatment. The first three groups were hypothesized to constitute a Principal Response. To validate these groupings, we constructed protein-protein interaction (PPI) networks using the Human Protein Atlas and STRING. The main contribution of this work is the implementation of second-order connections restricted to those made via DEGs, ensuring connectivity reflects condition-specific responses rather than generic hubs. We show applying our DEG filter to second-order connections can change the conclusions drawn from the same selected gene sets. Support Group genes also showed strong interconnectivity and overlap with housekeeping genes. STRING confirmed PPI enrichment but produced less stable results. By emphasizing DEG-restricted second-order protein interactions, we address limitations of context-free enrichment methods and strengthen biological evaluation of computational models of differential gene expression. Our framework is designed to support the prioritization of subsets of DEGs for experimental follow up but not to predict exactly how an environmental input produces a physiological output.

**NEW & NOTEWORTHY:** Computational models aim to identify differentially expressed genes (DEGs) that respond to environmental perturbations. Models usually select genes thought to be of elevated biological importance. We present a framework for supporting the mechanistic plausibility of DEGs using proteomic data. This simple but generalizable algorithm develops context-informed protein-protein interaction networks, known to reflect genetic epistatic networks, providing a tool to test the biological plausibility of model-selected genes in forming adaptive response mechanisms prior to experimental validation.

## INTRODUCTION

Computational models are increasingly deployed to identify genes that are differentially expressed in response to various conditions. Standard approaches for selecting differentially expressed genes (DEGs) typically rely on statistical comparisons of transcript counts before and after environmental effects, or between phenotypes. For example, the *limma* package in R first fits a linear model before running statistical tests to identify likely DEGs based on expression magnitude and variance modeling [1]. Other commonly used methods similarly use moderated t-statistics that compare geometric means of expression counts [2]. See [3] for a comparative review of the statistical tools for finding DEGs. These methods aim to distinguish genuine changes in gene expression associated with environmental exposures or phenotypic differences from the inherently noisy background of RNA-seq data. They are effective when replicate numbers are sufficient and fold-change differences are the primary outcome of interest. However, a novel approach has been developed for studies with few replicates or when inter-individual variability itself may have biological significance.

The method described in Gonzalez et al. (2026), uses other approaches to identify and classify DEGs, specifically those that maintain homeostasis in mammals in response to changing temperature [4]. This model is based not on mean fold-change, but on changes in inter-individual variability of transcript abundance before and after treatment. Using RNA-seq data from five human skin fibroblast samples exposed to decreased (32°C) or increased (41°C) temperature compared to baseline (37°C), the model defined DEGs as genes whose activity (in absolute value) after treatment exceeded baseline activity. For a detailed explanation of the cell culture methods, experimental treatments, and RNA-seq processing, see sections 2.1-2.4 in [4]. Briefly, the model compared baseline inter-individual variation in RNA transcript counts to post-treatment variation to identify genes most likely responsible for the principal response of the species. Of the 9,825 candidate human genes, 2,810 were identified to be DEGs in the 32° treatment and 2,679 in the 41° treatment. For a detailed explanation of how DEGs were identified, see definition 4.2 in [4]; the goal of the present study is not to describe or compare DEG-selection models, but rather to evaluate connectivity within the selected DEG subsets and determine whether restricting shared interaction intermediaries to condition-specific DEGs change that assessment.

Among the DEGs, the Gonzalez Model categorized extreme-response genes into four groups based on their positions in a before-versus-after inter-individual variability plot (Figure 1) [4]. Genes located at the extreme corners of this distribution were categorized as Key-Response (g1), Treatment-Specific (g2), Support (g3), and Noisy (g4). The Key-Response, Treatment-Specific, and Noisy groups are proposed to comprise the Principal Response Genes responsible for driving the characteristic species-level response to temperature change. Support Genes were thought to be well-regulated like the Principal Response genes, but also likely to be involved in many processes for many species across conditions. Because housekeeping (HK) genes are classically defined as genes that are consistently expressed across different cell types and conditions to support essential cellular functions such as metabolism, transcription, and maintenance, we suspected that many Support Genes may overlap with HK genes. Housekeeping genes and the proteins they encode are also known to exhibit high degree (i.e., a high number of known interactions) in protein-protein interaction networks because their central cellular roles require participation in numerous molecular pathways and physical interactions, and because their genes, being evolutionarily ancient, have had more time to accumulate such interactions. In addition to evaluating condition-informed network connectivity, we test whether housekeeping genes are disproportionately assigned to Support Group/g3. This group is defined by high inter-individual transcription variability both before and after treatment. We test both preferential allocation to g3 relative to the other model groups and enrichment of housekeeping genes in the group relative to the full candidate-gene background. See Support Group/G3 Genes in the Results and Discussion for more.

**Figure 1.**
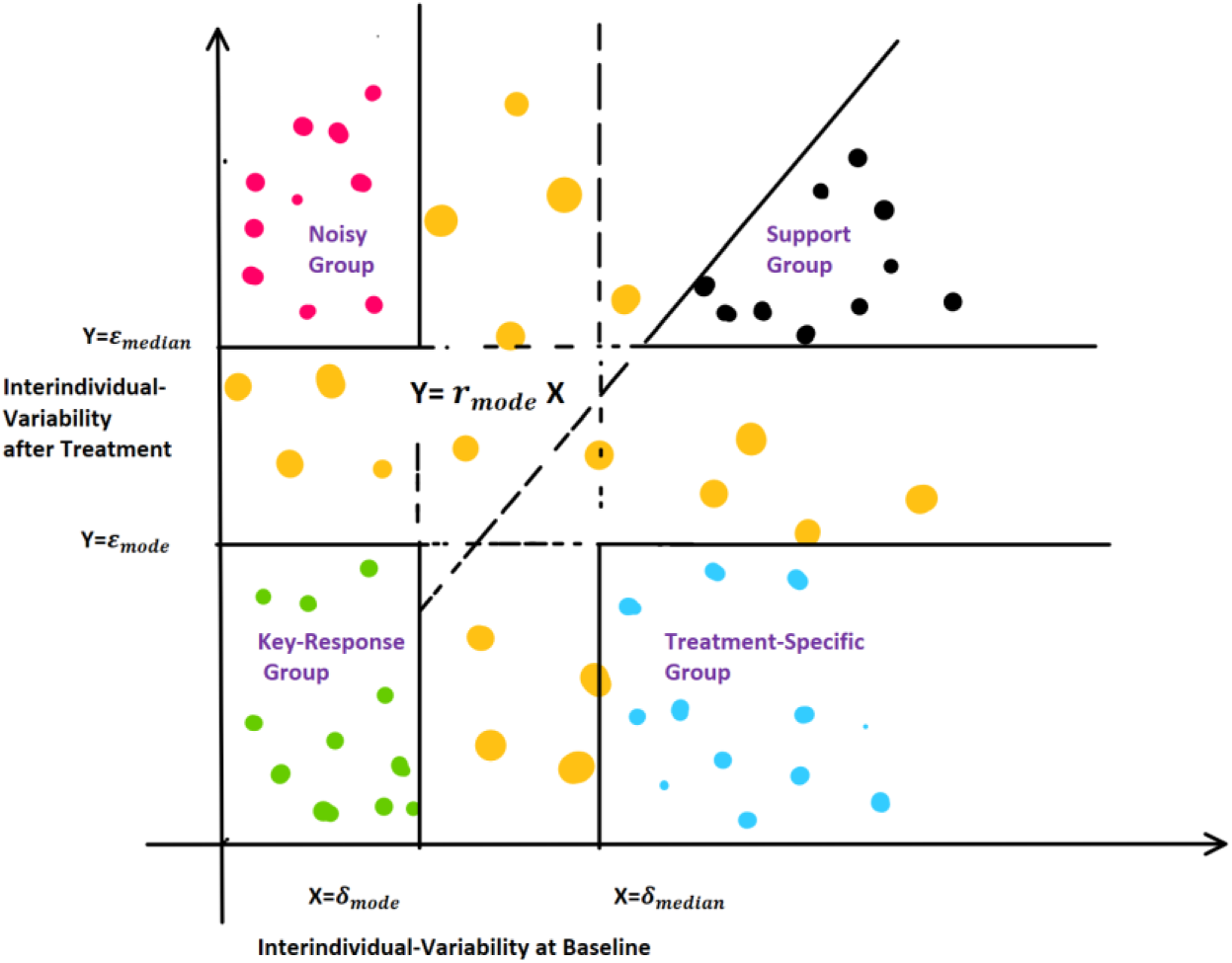
Graphical representation of the groups from the Gonzalez Model evaluated in this study [4]. Certain differentially expressed genes (DEGs) were classified according to their relative inter-individual variability at baseline and after treatment. Genes located at the extremes of the distribution depicted were categorized into four groups: Key-Response (g1), Treatment-Specific (g2), Support (g3), and Noisy (g4). Key-Response, Treatment-Specific, and Noisy groups are considered to make up the Principal Response. Reproduced from Gonzalez, J., Genereux, D. P., Crouse, K., et al., “New approaches to discovering epigenetic rules of homeostasis in diverse mammal species,” BMC Genomics 27, 478 (2026), Fig. 3, Springer Nature, with permission from Springer Nature Customer Service Centre GmbH. RightsLink License No. 6276020476742. No changes were made.

A major challenge for novel DEG-selection models is validation: how can we determine whether the identified genes truly participate in a coherent biological mechanism rather than reflect statistical noise? Here, we present a framework that gathers and analyzes functional proteomic data found in the literature to test whether a given gene set is likely to contribute to a coherent biological mechanism and whether the application of a DEG-filter to second-order intermediaries (i.e., condition-derived information) can change that assessment. The purpose of this is to serve as a preliminary indication that the gene set has literature-support, however further experimental analysis would need to be conducted to validate an entire network. Experimental validation is especially necessary when there is a paucity of published data for the tested organism’s proteome. This may include simultaneous knockdown studies, pairwise epistasis or double-perturbation analysis, and Perturb-seq approaches that combine CRISPR knockout with single-cell transcriptomics [5–7]. Notably, Castellanos-Girouard et al. (2026) found that pairwise double-mutant analysis revealed genetic interaction patterns that strongly correspond with protein-protein interaction networks, supporting the use of proteomic data to validate predicted gene relationships, as we present here [8]. Though preliminary in nature, computational methods, such as analysis of online databases, provide a less resource-intensive and more practical approach when studying large gene sets or multiple treatments and conditions before pursuing experimental validation.

For genes to act in a coordinated response to an environmental change, such as extreme heat or cold, we assume that their protein products must associate in a significant way (i.e., their proteins would form a network somewhat resembling a mechanism to promote homeostasis). Protein-protein interaction data could therefore be used to surmise whether software-identified DEGs are likely to carry out the function which the model studied. In other words, protein-protein interaction data could serve as a useful intermediate validation strategy between purely computational gene network prediction and full experimental validation. A central obstacle in current validation techniques is that protein-protein interaction (PPI) databases are not necessarily comprehensive or specific to the cellular contexts of interest (such as cell type and treatment condition). Existing enrichment tools, such as STRING-db, often rely on broad background interaction maps and may be biased toward highly studied genes. Specifically, second-order interactions can be inflated by generic hub proteins, making PPI enrichment alone limited in determining whether a proposed DEG subset reflects a mechanism of interest.

To address these challenges, we developed a context-specific protein interaction network validation framework that integrates biological literature and DEG-specific second-order gene interactions, providing a more targeted alternative to enrichment tools such as those found on STRING-db.org. We use gene sets produced by the Gonzalez model [4] and limma as test cases to evaluate whether restricting 2^nd^ order protein interactions to condition-specific DEGs preferentially preserved gene connectivity relative to random (degree-matched) gene sets.

Our framework could generally be used to validate the biological significance of any set of differentially expressed genes (of a well-studied organism) that includes a specific subset, which the researchers determined to be of particular interest. Whether due to variability patterns, mean-fold change after treatment, or enrichment annotations, studies of differentially expressed genes often home in on a narrower list of genes among the many. Our method can test whether this subset of genes’ protein products are known to be more interconnected than would be expected by chance under degree-matched controls. From this information, we gain insight into the biological evidence and trustworthiness of DEG models before validating them experimentally.

In the following, we apply this framework to Principal Response and Support gene sets from the Gonzalez Model [4] and to limma-selected subsets of matching size. We ask whether these sets form larger LCCs than approximately degree-matched random sets, and whether their LCCs lose fewer members than random sets when shared intermediaries are restricted to DEGs. Finally, we characterize housekeeping allocation and interaction degree in G3.

### Algorithm 1. Pseudocode describing construction of first-afirst-and DEG-rrestricted-second-order-second-order PPI Network.

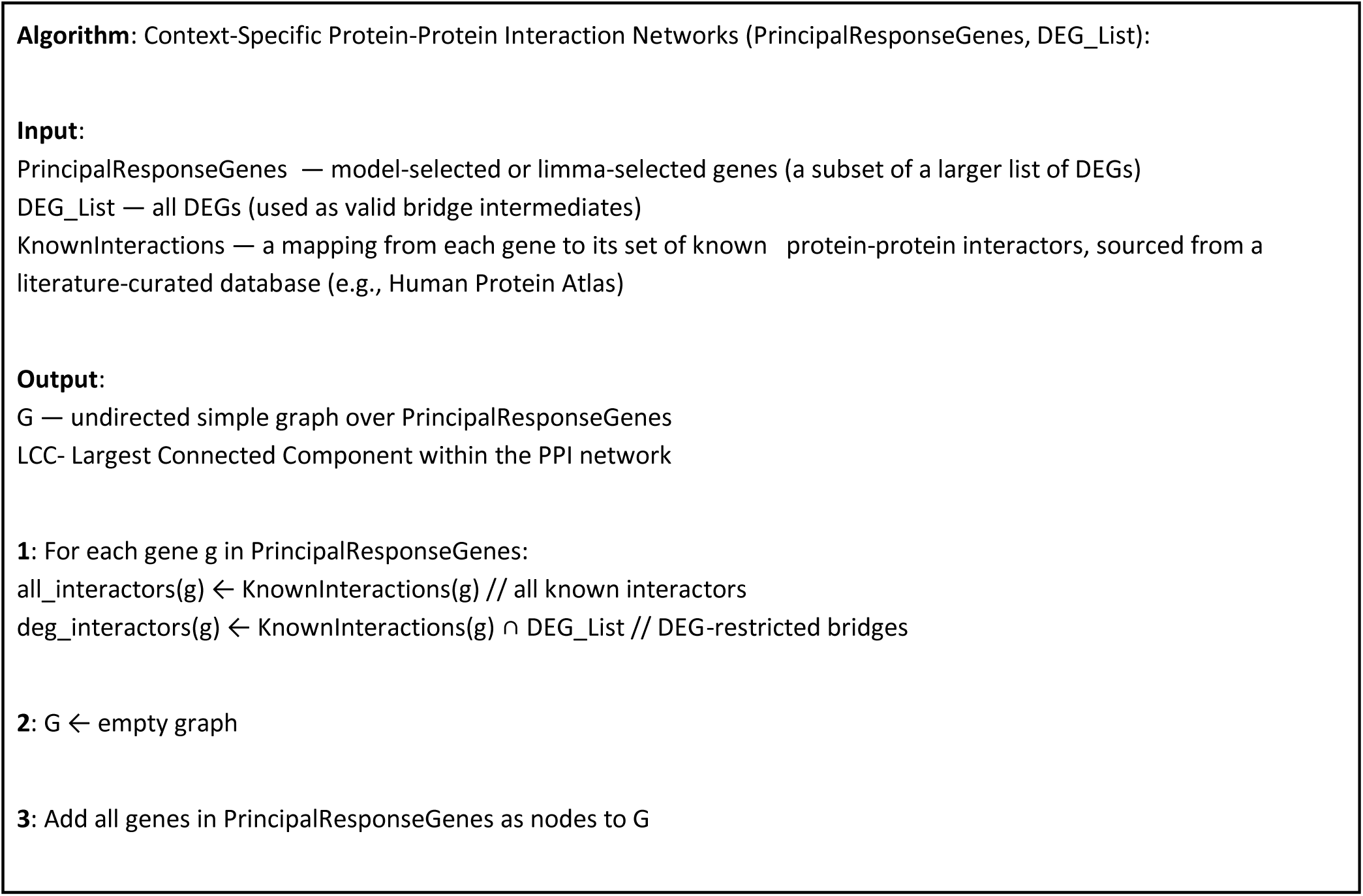
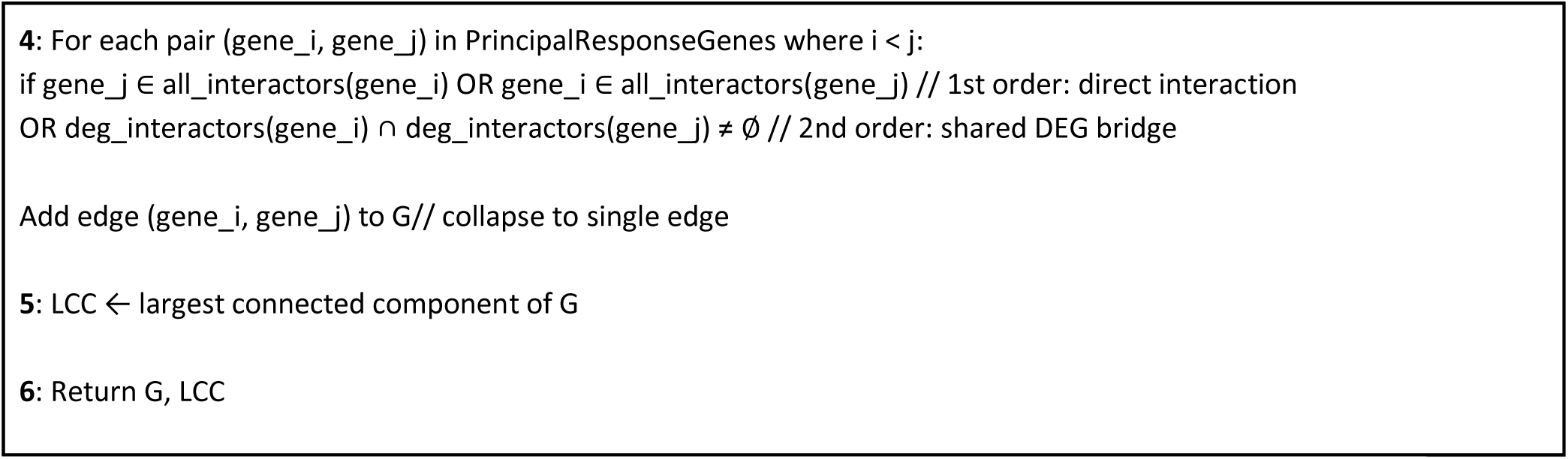

## MATERIALS AND METHODS

### Gene Sets and Study Design

The main contribution of this work is the restriction of shared intermediaries that generate second-order links in protein-protein interaction (PPI) networks to differentially expressed genes (DEGs) to make network assessments more condition-specific. To test how this changes PPI network connectivity, we use gene sets selected by the Gonzalez Model [4] and limma [1]. Our first comparison asks whether the selected gene sets lose fewer members from their largest connected components (LCCs) after applying our framework’s DEG restriction compared to approximately degree-matched random gene sets. A smaller decrease in LCC size than expected would suggest that the selected genes are connected through proteins whose encoding genes also respond to the temperature treatment (i.e., DEGs), rather than depending as heavily on the broader genome to connect them.

For each temperature condition (32° and 41°), two networks were built: one from the genes found in the Key-Response, Treatment-Specific, and Noisy groups (Principal Response Genes denoted as G1, G2, and G4, respectively) and a second from the Support Group (denoted as G3) genes only. We then used sets of limma-identified DEGs that were produced from the same RNA-seq data for each temperature condition. We selected the same number of highest-ranked limma genes as Principal Response genes (and separately, Support Group genes; Supplementary Table S5-S8) at each temperature.

### First and Second Order Connections (Network Edges)

First order connection: A direct model-gene-to-model-gene protein-protein interaction. Protein interactions were obtained from the Human Protein Atlas (PA) “Interaction” section, which aggregates data from four major sources: IntAct (high and medium confidence physical associations), BioGRID (multivalidated physical interactions), BioPlex (interactions with >75% probability), and OpenCell (significant physical interactions) [9]. We included any known interaction reported in at least one of these four major sources.

Second order connection: One in which two model-identified genes are connected via a literature-validated interaction with the same differentially expressed gene (as defined by the Gonzalez Model). In other words, a bridge between model genes A and B would be formed if they shared an interaction with the same DEG. For all selected sets at a given temperature, eligible bridges came from the same full condition-specific DEG list, rather than only the selected subset.

### Network Construction

Public resources found on the Human Protein Atlas (https://www.proteinatlas.org/) were used to identify the known interactions among genes found in the model-produced sets. A Python (version 3.12) web scraping script employing the requests library was used to obtain these known interactions. From there, custom Python functions were used to construct the networks outlined below and calculate LCC sizes and edge counts, with NumPy used for degree-bin construction (described below) and numerical calculations. Protein-protein interaction network construction included both first-order direct gene connections and second-order connections (see Figure 2). As stated, second-order interactions were only considered if the intermediate (bridge) gene was itself also a DEG.

**Figure 2.**
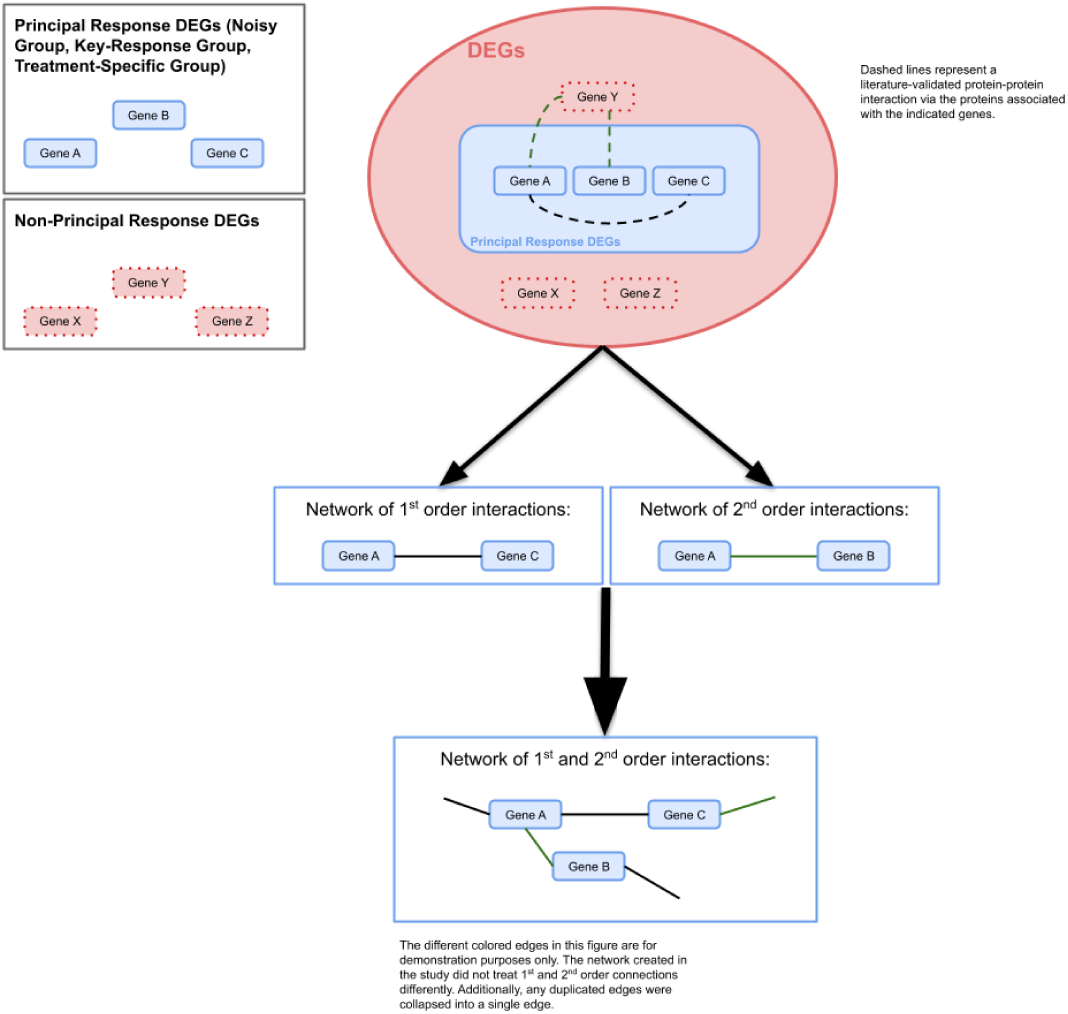
Graphical Explanation of the networks created in this study to validate the tested gene sets. 1^st^ order interactions are counted when two Principal Response DEGs are known to interact in the database, whereas 2^nd^ order interactions are counted when two Principal Response DEGs are known to interact in the database through a shared DEG intermediate.

Largest connected component (LCC) size was used as the primary network measure for testing the model, as we assumed it better reflects mechanistic coherence than edge counts. The LCC is a measure of the largest subset of genes that are mutually connected through known protein-protein interactions found in the databases. Each gene’s corresponding protein constitutes a node in the network, linked to others by edges representing the known interactions. Edges were added according to the two connection criteria described above.

The network test statistic used should reflect the goal of the model being tested. In our case, the Gonzalez Model attempts to precisely find the mechanism of the highest importance (referred to in [4] as the group of genetic drivers which make up the high order response of the species to the treatment). If the model truly identifies the genes responsible for the principal temperature response mechanism (as it sought to do), then those genes should not appear as isolated entities (see Figure 3). Instead, they would ideally form a cohesive mechanism, and evidence of such a mechanism is the presence of consistent and thorough protein–protein interactions that make for a large LCC.

**Figure 3.**
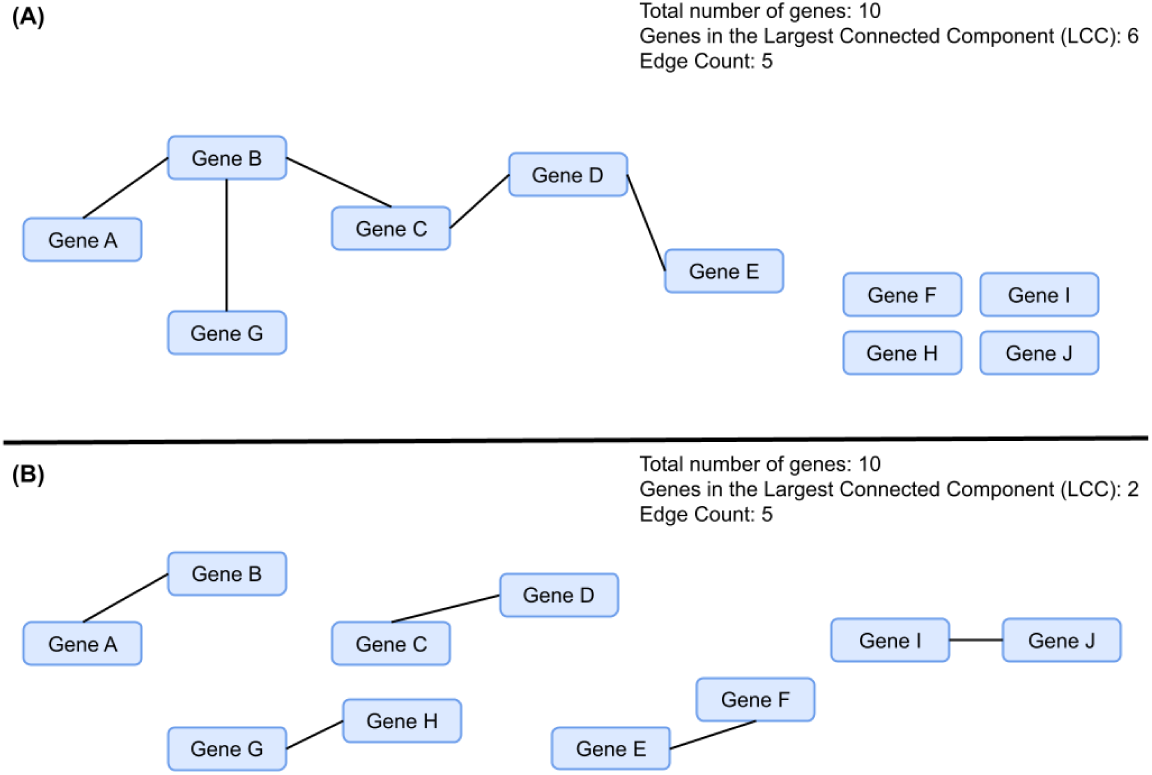
Gene networks consist of the same number of genes with different make ups. We posit that (A) is more reflective of a driving biological mechanism to the temperature response than (B), despite their equal edge counts.

When considering the networks formed by both first and second order interactions, edges were counted identically, regardless of the type (first or second order). The algorithm also collapsed multiple interactions between the same two genes and formed a simple (no parallel edges), undirected graph (Figure 2). The networks constructed for this study did not distinguish upregulated from downregulated genes. Edges were not assigned activating or inhibitory effects. Our framework could, however, be adapted to incorporate these distinctions (where supporting data are available) if researchers have more specific biological questions beyond the scope of the present study

### Degree-Matched Random Sampling

Degree-matched random sampling (using 20 bins of genes ordered by number of known interactions in Protein Atlas) provided null distributions against which observed connectivity could be compared. To create the null comparison, genes from the full background list of candidate genes studied (this included DEGs and non-DEGs) were binned by known degree. Random sets of genes were then selected from these genes, with the number of genes pulled from each bin matching the degree-distribution of the sample gene set. The candidate list of 9825 genes was used as a background as opposed to the whole genome to control for bias: In the likely case that our 9825 candidate genes are more well-studied, and therefore have more known protein-protein interactions, then we should expect the DEGs’ PPI networks to have an inflated connectedness if compared to the entire genome. Put simply, approximately degree-matched randomizations are required to correct for how some genes are more well-studied than others [10]. PPI networks are known to exhibit strong degree-dependent connectivity, where proteins that are generally highly connected are disproportionately likely to interact with each other [11]. 1,000 randomized gene sets were generated using this degree-approximation sampling procedure, and networks were created for each randomized set using the same first-order and DEG-restricted second-order edge rules applied to the observed gene set.

### Connectivity Retention After DEG Restriction

To isolate the effect of our framework’s restriction on 2nd order intermediaries to only DEGs, we constructed unrestricted networks allowing any recorded interactor to form a second-order edge. The selected genes (nodes), degree-matching, and 1,000 random gene sets were identical to those used for the restricted networks. Each observed and random set therefore had paired restricted and unrestricted results.

We tested whether the model-selected gene sets (which ostensibly should form networks more so structured by condition-informed intermediaries) preferentially retained LCC members when shared intermediaries were restricted to DEGs, relative to the losses observed in degree-matched random sets. For each set, we calculated Δ LCC = LCC (unrestricted bridges) – LCC (restricted). We calculated empirical values using p = (a + 1)/(1,000 + 1), where a was the number of random ΔLCC values ≤ observed ΔLCC. Smaller Δ LCC values indicate greater retention of LCC members relative to the random sets when condition information is applied. A small p value would indicate that the selected set lost unusually few LCC members after implementing our DEG-restriction on 2nd order interactions.

### LCC Size and Edge Count Enrichment

We also tested whether each restricted and unrestricted network had an unusually large LCC or an unusually high number of edges within its LCC relative to the corresponding random networks.

The 1,000 randomized networks were used to estimate the expected null distribution of the LCC size and LCC edge count, against which the observed network (both before and after applying our DEG-filter to 2^nd^ order intermediaries) statistics were compared using upper-tail empirical rank tests. Upper-tail empirical p-values were calculated as p = (b + 1)/(1,000 + 1), where b was the number of random networks whose LCC size (or LCC edge count) equaled or exceeded the observed value.

### Housekeeping Gene Analyses

As previously mentioned, we suspected that group 3/Support Group genes might be enriched for Housekeeping Genes. To test this, in addition to performing the network analysis described above for solely these genes, we also cross-referenced a known housekeeping gene database. We used the Housekeeping and Reference Transcript Atlas (version 1.0), a web database which mined high-quality human RNA-seq data sets [14].

To test preferential allocation of housekeeping genes within the Gonzalez Model, we compared G3 with the union of G1, G2, and G4 using a one-sided Fisher’s exact test on housekeeping vs non-housekeeping counts. We report both the share of all model-selected genes and the share of model-selected housekeeping genes found in G3 for each temperature condition. Separately, we tested G3 housekeeping enrichment against the candidate gene background using one-sided Fisher’s exact tests after G3 genes were excluded from the comparison group.

For each temperature, we compared G3’s median degree with medians from 1,000 random gene sets of the same size, sampled from the 9,825-gene background. Unlike the network comparisons, these random sets were not degree-matched because degree itself was the property being tested. We calculated upper-tail empirical p-values, using the formula p = (c + 1)/(1,000 + 1), where c was the number of random gene sets whose median degree equaled or exceeded that of the observed G3. We repeated this analysis after removing housekeeping genes from both G3 and the background to determine whether elevated degree was also evident among G3’s non-housekeeping members.

We calculated the mean, median, and standard deviations of the number of known interactions for the Principal Response Group, Support Group, entire DEG set, and background gene set for each temperature condition.

### STRING-db Comparison

STRING, a database of known and predicted protein-protein interactions, was also used to generate a PPI enrichment p-value for the gene sets. The web database version 12.0 was used [12, 13]. For the analysis, the whole genome was used as the statistical background (which is default to the website) since our background list of 9825 was too large to upload. Default settings were used: full STRING network (the edges indicate both functional and physical protein associations) composed of interactions from text mining, experiments, databases, co-expression, neighborhood, gene fusion, and co-occurrence; medium confidence (0.400) minimum required interaction score.

### Software and Code Availability

Python version 3.12 and its packages requests, NumPy, and SciPy were used for all final analyses. Generative artificial intelligence tools, namely OpenAI’s ChatGPT, were used in the development of the Python scripts used in this study. The source code used is available at https://github.com/BradleyFrishman/Context-Specific-DEG-PPI-Network-Builder.

## RESULTS

### Connectivity Retention After DEG Restriction

The Gonzalez Model’s Principal Response genes retained more members of their LCC compared to random gene sets after applying our DEG filter to second order bridges (statistically significant at 32°, not at 41°) (**Table 1**).

**Table 1.**
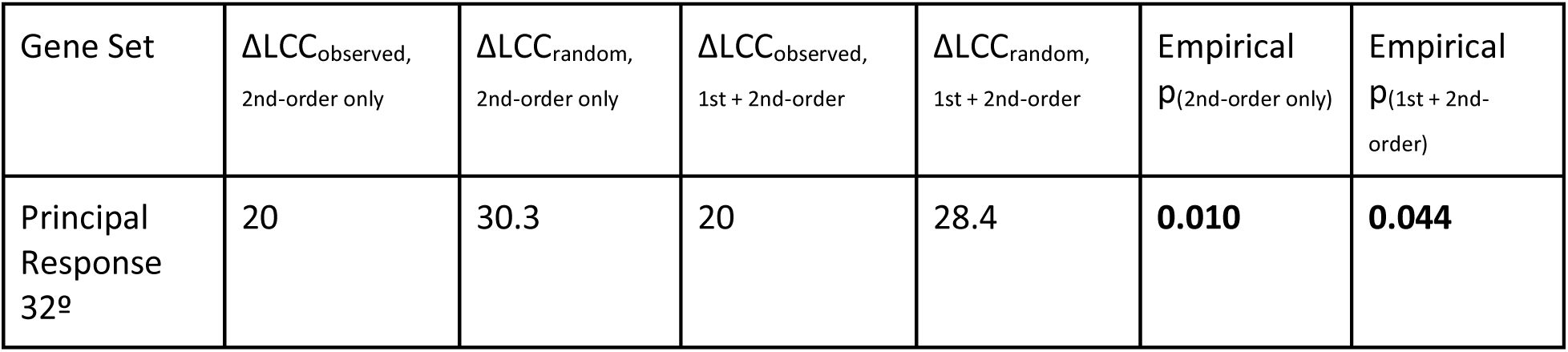
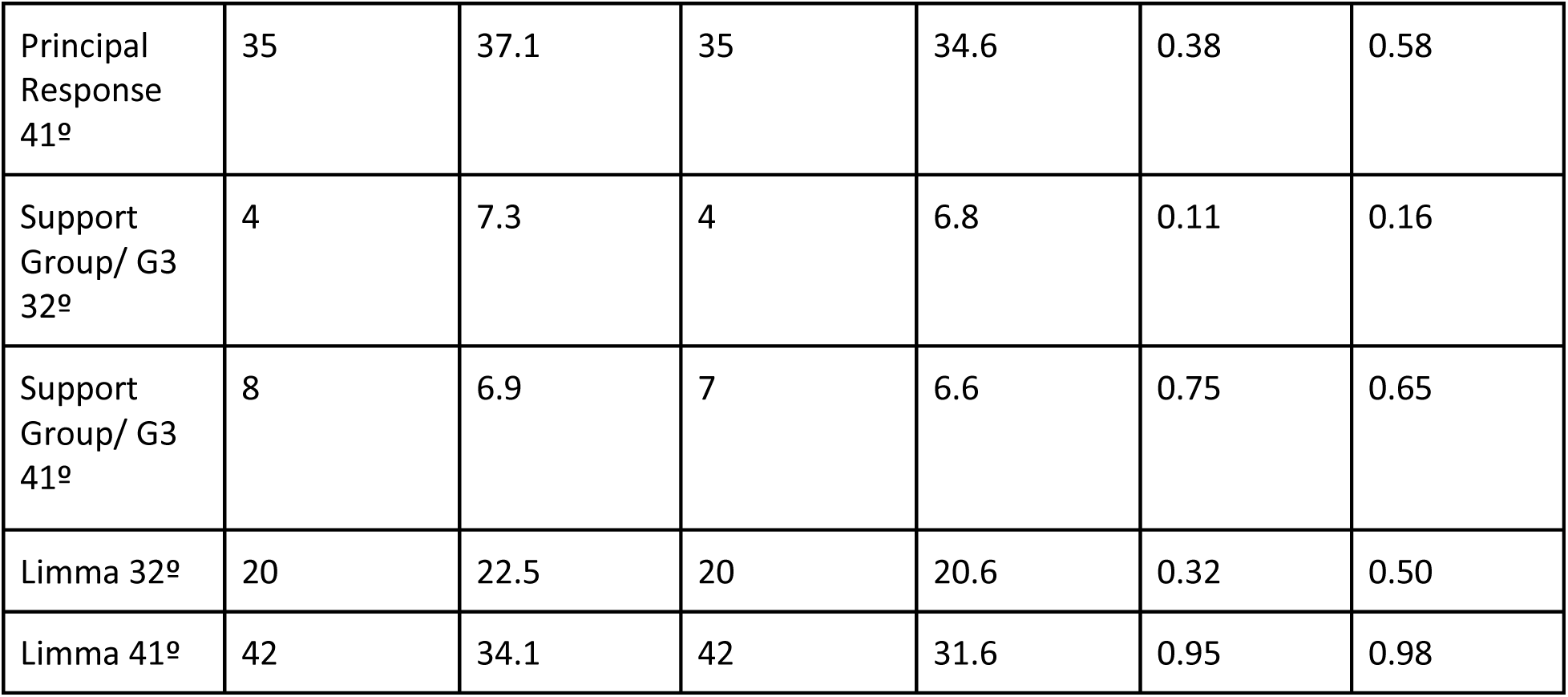
Results from the given network of genes that the Gonzalez Model identified to be a part of the Principal Temperature Response (Principal Response Group: groups 1,2, and 4), a same sized set of top-ranked limma-selected DEGs, or Support Group (Group 3). Unadjusted p-values generated from a one-sided empirical rank test.

| Gene Set | $\Delta LCC_{\text{observed, 2nd-order only}}$ | $\Delta LCC_{\text{random, 2nd-order only}}$ | $\Delta LCC_{\text{observed, 1st + 2nd-order}}$ | $\Delta LCC_{\text{random, 1st + 2nd-order}}$ | Empirical $p_{(2\text{nd-order only})}$ | Empirical $p_{(1\text{st} + 2\text{nd-order})}$ |
| --- | --- | --- | --- | --- | --- | --- |
| Principal Response 32<sup>o</sup> | 20 | 30.3 | 20 | 28.4 | **0.010** | **0.044** |
| Principal Response 41° | 35 | 37.1 | 35 | 34.6 | 0.38 | 0.58 |
| Support Group/ G3 32° | 4 | 7.3 | 4 | 6.8 | 0.11 | 0.16 |
| Support Group/ G3 41° | 8 | 6.9 | 7 | 6.6 | 0.75 | 0.65 |
| Limma 32° | 20 | 22.5 | 20 | 20.6 | 0.32 | 0.50 |
| Limma 41° | 42 | 34.1 | 42 | 31.6 | 0.95 | 0.98 |

The paired ΔLCC analysis tested whether the decrease under DEG restriction was smaller than expected from random sets. At 32°C, the Gonzalez-Model Principal Response second-order-only LCC decreased from 153 to 133 genes, a loss of 20, compared with an average loss of 30.3 in random networks. This loss was significantly smaller than expected (empirical p = 0.0100). The combined-network LCC similarly lost 20 genes, compared with 28.4 on average in random networks (p = 0.0440).

At 41°C, the Principal Response second-order-only LCC decreased from 175 to 140 genes, a smaller loss of 35 compared with 37.1 on average in random networks; this difference, however, was not significant (p = 0.3846). The combined-network comparison was also not significant (p = 0.5794).

For an equal number of top ranked limma-identified differentially expressed genes from the same RNA-seq data at 32° and 41°, the gene networks did not show significantly smaller than expected decreases in their LCC after applying our DEG filter to second-order bridges. The same is true for the Group 3/Support Group Genes.

### LCC Size

The Gonzalez Model’s Principal Response genes formed larger than expected LCCs at 32° but not 41° after applying the DEG filter to second-order intermediaries.

At 32°, Principal Response Group genes formed a significantly larger LCC in its PPI network than expected by chance (**Table 2**). Of the 182 genes, 167 were included in the network analyses after excluding those with no known interactions in the database. The DEG-restricted second-order only network formed an LCC of 133 genes, compared with an average of 123.4 in random networks (empirical p = 0.0250). This significance was not present for the unrestricted second-order only network (empirical p = 0.70; Supplementary Information Table S4). The combined first-and second-order LCC contained 134 genes and was also significantly larger than expected (random mean = 125.5, p = 0.0480).

**Table 2.** Results from the given network of genes that the Gonzalez Model identified to be a part of the Principal Temperature Response (Principal Response Group: groups 1,2, and 4). Unadjusted p-values generated from a one-sided empirical rank test.

| Gene Set, Network | LCC size | Null mean LCC size | LCC size p |
| --- | --- | --- | --- |
| Principal Response 32°, 2nd-order only | 133 | 123.4 | **0.025** |
| Principal Response 32°, 1st + 2nd-order | 134 | 125.5 | **0.048** |
| Principal Response 41°, 2nd-order only | 140 | 133.2 | **0.12** |
| Principal Response 41°, 1st + 2nd-order | 141 | 135.8 | **0.19** |

At 41°, 186 out of 204 of the Principal Response genes were included in the network analyses after excluding those with no known interactions in the database. The gene set did not form a significantly larger LCC compared to the 1,000 random degree-approximated gene sets (p = 0.1229 and 0.1918 for the second-order only and combined networks, respectively) (**Table 2**). The unrestricted LCC for the second-order interaction-only network was borderline significantly large (p=0.049), however.

At 32°, a Principal-Response-sized subset of *limma*-identified differentially expressed genes formed a larger LCC than the degree-matched random expectation. For both the second-order only and the combined first and second-order networks, the genes assembled into a LCC of 149 genes compared to random means of 138.8 (p = 0.0070) for second order-only and 140.8 for combined networks (p = 0.0280). At 41°, PPI networks formed by *limma* genes were not significantly large. Both LCCs (solely second order and first and second order combined) contained 142 genes, fewer than the random means of 143.1 and 145.7 genes (p = 0.6234 and 0.7872) (**Table 3**).

**Table 3.** Results from the limma subset of genes matched to the number of genes that the Gonzalez Model identified to be a part of the Principal Temperature Response (Principal Response Group: groups 1,2, and 4). Unadjusted p-values generated from a one-sided empirical rank test.

| Gene Set, Network | LCC size | Null mean LCC size | LCC size p |
| --- | --- | --- | --- |
| 32°, 2nd-order only | 149 | 138.8 | **0.0070** |
| 32°, 1st + 2nd-order | 149 | 140.8 | **0.028** |
| 41°, 2nd-order only | 142 | 143.1 | 0.6234 |
| 41°, 1st + 2nd-order | 142 | 145.7 | 0.7872 |

### Support Group/G3 Genes

The Support Group genes, Group 3 (G3), showed evidence of increased connectivity and overlap with known housekeeping (HK) genes. After removing genes with no known interactions in the database, G3 gene count falls from 67 to 62 at 32°C and from 53 to 46 at 41°C. These genes will be used for the purpose of the following network analyses; housekeeping and degree analyses use all 67 and 53 genes.

First, using the same network-construction and null comparison applied to other groups, the LCCs of the G3 gene set were found to be significantly larger than expected for the 32° condition but not the 41° (**Table 4**). Furthermore, unlike the Principal Response Genes (union of G1, G2, and G4) network, the 32° G3 networks had significantly larger than expected edge counts in their LCCs (not observed for 41° DEG-restricted set). However, the G3 genes’ PPI networks did not produce significantly smaller-than-expected changes in their LCC after applying our DEG filter to second-order bridges.

**Table 4.** Results from the Gonzalez Model Group 3/Support Group genes. Unadjusted p-values generated from a one-sided empirical rank test.

| Gene Set, Network | LCC size | Null mean LCC size | Observed LCC edge count | Null LCC edge count mean | LCC size p | Edge Count p |
| --- | --- | --- | --- | --- | --- | --- |
| 32°, 2nd-order only | 57 | 50.0 | 469 | 308.2 | **0.0030** | **0.0010** |
| 32°, 1st + 2nd-order | 57 | 50.6 | 476 | 312.0 | **0.0040** | **0.0010** |
| 41°, 2nd-order only | 36 | 35.0 | 165 | 143.0 | 0.4316 | 0.1998 |
| 41°, 1st + 2nd-order | 37 | 35.4 | 168 | 144.8 | 0.3257 | 0.1888 |

A higher proportion of G3 genes overlapped with an established housekeeping gene database than observed for the Principal Response gene sets (<12%) and the overall background prevalence (21.77% of all 9,825 genes). Specifically, 18 of 67 G3 genes at 32°C (26.9%) and 18 of 53 G3 genes at 41°C (34.0%) were found in the housekeeping database. Compared with the background, these represented 1.23-fold and 1.56-fold enrichment, respectively, although only the 41°C comparison was statistically significant (32°C: p = 0.19; 41°C: p = 0.028) (**Table 5**).

**Table 5.** Results from housekeeping (HK) gene overlap and allocation analyses on Support Group/G3 genes.

| Measure | 32° | 41° |
| --- | --- | --- |
| HK overlap within G3 | (18 / 67) = 26.87% | (18 / 53) = 33.96% |
| HK overlap within background | (2,139 / 9,825) = 21.77% | (2,139 / 9,825) = 21.77% |
| Fold enrichment in G3 versus full background | 1.23 | 1.56 |
| Fisher p versus background (excluding G3) | 0.19 | **0.028** |
| G3 share of model-selected genes | (67 / 249) = 26.91% | (53 / 257) = 20.62% |
| Share of model-selected HK genes in G3 | (18 / 38) = 47.37% | (18 / 42) = 42.86% |
| Fisher p for G3 versus G1+2+4 | **0.0027** | **0.00027** |

Housekeeping genes were disproportionately assigned to G3 instead of other groups in the Gonzalez model at both temperatures. Another way we examined the association between housekeeping genes and the Support Group genes was by testing whether housekeeping genes assigned to any of the four groups were disproportionately found in G3. At 32°C, G3 accounted for 26.9% of all model-selected genes but contained 47.4% of their housekeeping genes. At 41°C, G3 accounted for 20.6% of model-selected genes but contained 42.9% of their housekeeping genes. One-sided Fisher’s exact tests comparing G3 with the Principal Response groups were significant at both temperatures (32°C: p = 0.0027; 41°C: p = 0.00027) (**Table 5**).

G3 genes tended to have a higher number of known interactions in the database (Supplementary Figure S2, Supplementary Table S3). Their median number of known interactions was 22 at both temperatures, compared with 13 in the background. Comparisons with 1,000 randomly sampled gene sets of the same size, without degree matching, supported this difference at both temperatures (32°C: p = 0.0160; 41°C: p = 0.0260; Supplementary Table S1). This suggests that the subgroup may include more hub-like genes, meaning genes whose proteins have many recorded interaction partners.

After removing known housekeeping genes from both G3 and the background, neither degree comparison remained statistically significant. The remaining 49 genes at 32°C had a median degree of 16, compared with 10 in the non-housekeeping background (p = 0.0729). The remaining 35 genes at 41°C had a median degree of 9, compared with 10 in the background (p = 0.7243; Supplementary Table S1). These results suggest that housekeeping members predominately contribute to G3’s higher degree, particularly at 41°C.

Notably, this grouping was created by identifying genes with high inter-individual variability both before and after treatment, completely independently of interaction degree or housekeeping annotation. Nevertheless, the selected genes had elevated median degree, and a disproportionate share of the model-selected housekeeping genes fell into G3. Within these data, selecting genes in this manner therefore identified a group with both high connectivity and substantial housekeeping status.

### STRING-db Comparison

STRING analysis supported that the union of G1, G2, and G4 model genes were enriched for protein-protein interactions with p-values of 5.86e-05 (32° condition) and 1.3e-08 (41° condition) (**Table 6**). These PPI enrichment p-values are generated by comparing the observed number of interactions with that expected using a random-graph model that accounts for protein degree [15]. This indicates that the input proteins have more known or predicted associations among themselves than what would be expected for a random set of proteins of the same size and degree distribution. Notably, the approximately degree-matched random gene sets showed widely varying PPI enrichment p-values, (frequently < 0.05) indicating significance for ostensibly random collections of genes.

**Table 6.**
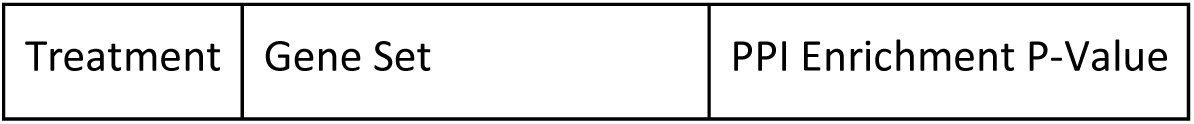
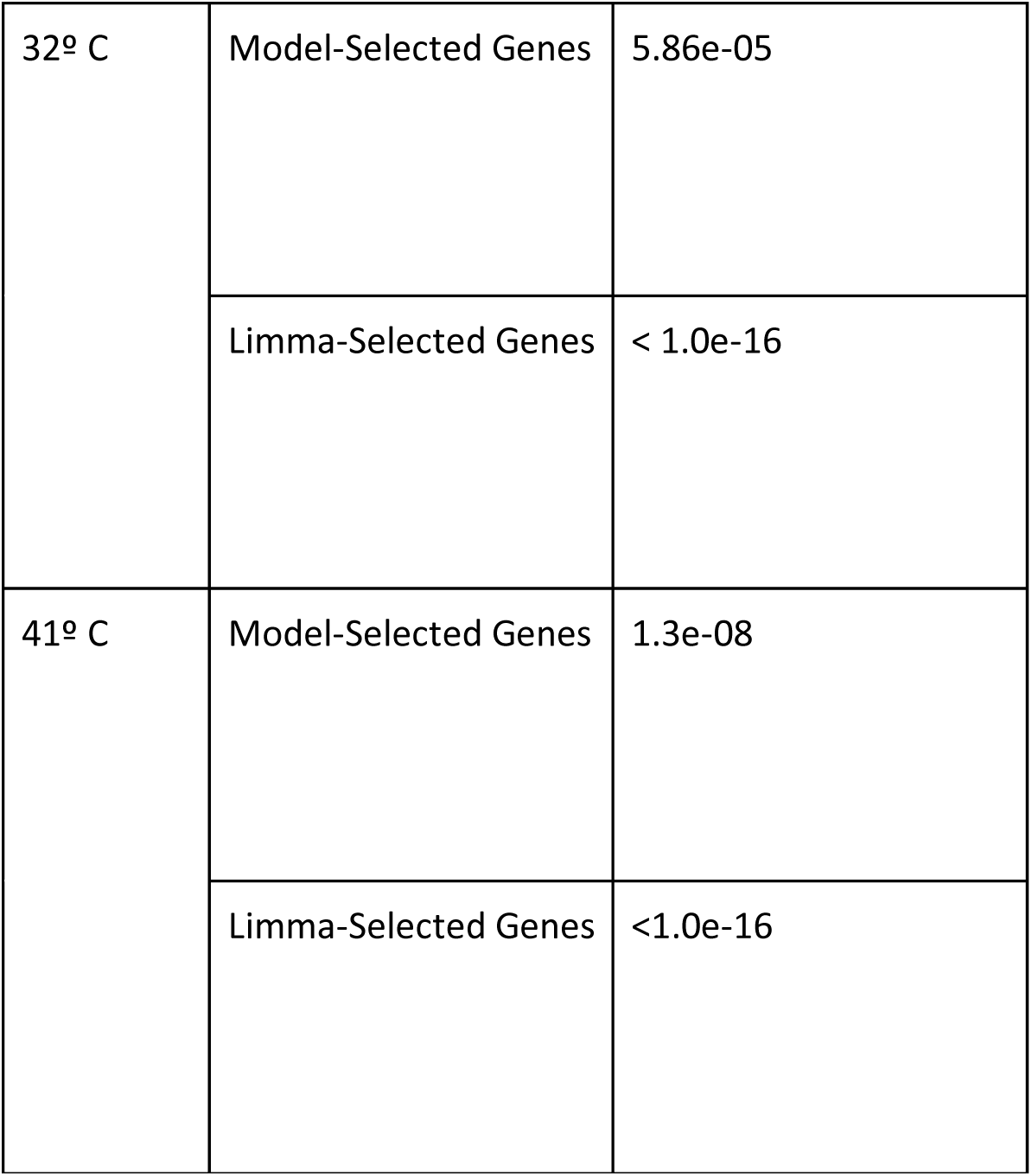
STRING-db Results. Protein-protein interaction enrichment analysis of the genes the Gonzalez Model identified to be a part of the Principal Temperature Response (Groups 1, 2, and 4) and the genes that limma selected from the same RNA-seq dataset. P-values are generated by comparing the observed number of interactions with that expected using a random-graph model that accounts for protein degree [15]

| Treatment | Gene Set | PPI Enrichment P-Value |
| --- | --- | --- |
| 32° C | Model-Selected Genes | 5.86e-05 |
|  | Limma-Selected Genes | < 1.0e-16 |
| 41° C | Model-Selected Genes | 1.3e-08 |
|  | Limma-Selected Genes | <1.0e-16 |

STRING analysis of the limma-identified differentially expressed genes from the same RNA-seq data found them to be enriched for protein-protein interactions as well (**Table 6**) for both temperature conditions. The p-values associated with the limma-selected genes are considerably smaller than the ones for the Gonzalez Model-selected genes; this may be because limma has been widely used for over a decade and has greatly influenced the direction of studies, further strengthening their association in the STRING database.

## DISCUSSION

The goal of this study was to evaluate a condition-informed protein-protein interaction framework that restricts shared intermediaries to identified DEGs. When compared to generic, unrestricted networks, applying our DEG filter to second-order connections can change the conclusions drawn from the same selected gene sets (Supplementary Information Table S4). Additionally, we find that at 32°C the Gonzalez Model’s Principal Response genes retained more LCC members after applying the filter compared to approximately degree-matched random sets. This statistically significant result was observed for both second order-only networks (p = 0.0100) and combined first- and second-order networks (p = 0.0440), supporting the idea that these model-selected genes preferentially retain their interconnectedness through DEG intermediaries. Whereas the randomly selected gene sets largest connected components relatively greatly lose members of their LCC after applying the DEG filter, model selected genes under cold-perturbation retained their second-order interactions.

For the 41°C treatment, Principal Response second-order-only network showed the same directional trend of a relatively small observed LCC loss compared to the random sets, but the difference was not significant (p > 0.05). Weaker support for PPI network connectivity at 41°C compared to the null model occurred frequently throughout our analyses. Whether this pattern reflects limitations of the framework, characteristics of the RNA-seq data experimental procedure, or biological differences between heat and cold responses remains unresolved.

At 32°C, this study shows that the Gonzalez Model identifies gene networks that have more evidence of being biologically interconnected than would occur by chance, underscoring the plausibility of its predictions and the abilities of our efficient framework. By restricting shared intermediaries in second-order connections to DEGs, our approach aims to enhance context specificity and account for data bias compared to traditional enrichment analyses. The observation that model-identified genes, along with the 32°C gene set identified by *limma*, assemble into strongly connected networks supports the novel computational framework’s effectiveness. If the selected genes take part in a species’ primary response to a particular treatment, they could act together as part of a mechanism. That single mechanism (or network) should be complete, ideally composed of most or all the selected genes. The literature-supported connectivity we observed is consistent with such a mechanism. The findings illustrate the utility of tailoring network validation frameworks to experimental conditions rather than relying on broad, genome-wide baselines.

Further support for considering shared protein interactors when evaluating gene sets comes from Besse et al. (2026). They found that pairs of protein-protein interactions sharing a binding partner (i.e., A-B and B-C, which would constitute a second-degree connection between A and C in our framework) shared more protein-interaction quantitative trait loci (piQTLs) than pairs without a shared binding partner. This finding emphasizes the biological importance of indirect protein-protein connections in linking genetic variation to phenotypic changes, which our framework further filters to include condition-informed specificity [16].

Principal Response Group-sized gene sets identified by *limma*, which is considered standard DEG software, from the same RNA-seq raw data also produced a significantly larger LCC for the 32° condition but not for the 41° condition. At 41°C, LCCs were not significantly larger or smaller than expected. Unlike the Principal Response genes at 32°C, the limma subsets did not show significantly smaller than expected losses in LCC size after DEG restriction at either temperature. This may be due to the goal of the two DEG-identifying models. Whereas the Gonzalez Model aims to identify the genes that make up a driving mechanism underlying response to temperature change, limma may more generally seek to find any gene that is statistically up-or downregulated [1,4].

We assess that the results from STRING are less reliable than our own network building process for the following reasons: First, STRING’s enrichment p-values are calculated using the entire genome of up to 20,000 genes as the background (genes which can be used in the randomly generated/null comparison gene sets), whereas our own network analysis restricts comparison to the smaller list of 9,825 genes that were studied. Using a relevant and restricted universal gene set for randomization creates a more appropriate null distribution. Second, our approach only incorporates second-order connections that are made through differentially expressed genes, directly tying the network structure to the experimental condition of interest; STRING does not employ this specificity. Third, while more statistically involved than our calculations, STRING evaluates enrichment primarily by edge count, while we focus on the size of the largest connected component. We argue LCC can be a more biologically meaningful measure of a functional mechanism for our use case given the goal of the tested model. Similar reasoning underlies the protein interaction network analysis conducted by Hanes *et al.* (2023) in their study of stress response mechanisms in *Listeria monocytogenes* [17]. In that study, networks curated from STRING interaction data were refined using MCODE (Molecular Complex Detection) to emphasize highly interconnected clusters. This reflects the assumption that well-connected subnetworks are more biologically interpretable. MCODE identifies groups of proteins likely to form molecular complexes based on their interaction patterns, thereby using connectivity as a marker of functional coherence. Consistent with this framework, the most promising protein targets were found to be highly interconnected within a large network, implicitly treating membership in a well-connected component as evidence of mechanistic centrality. Rather than focusing on isolated or sporadic protein-protein interactions, the authors aimed to identify larger interaction components that could resemble functional biological systems.

The use of LCC as a primary network statistic is further supported by canonical stress-responses that are organized around direct physical protein-protein interactions. For example, bacterial proteostasis during heat stress is enacted by the chaperonin machinery centered on GroEL and its cofactor GroES, which function as a physically interacting subnetwork and require coordination with additional chaperones to mount an effective stress response [18]. Such systems illustrate how stress-response mechanisms manifest as cohesive interacting protein networks that work in concert to carry out their function.

While edge count quantifies the total number of protein-protein interactions within a gene set, it does not assess whether those interactions form a concentrated, integrated structure like the ones found to enact physical stress responses. The LCC can capture whether several genes are mutually reachable through interaction pathways, a prerequisite for coordinated function. Given that the goal of the Gonzalez Model is to identify functional temperature-response mechanisms rather than isolated interaction density, LCC was selected as the primary validation metric.

Housekeeping genes and the proteins they encode have been shown to exhibit high degree in protein-protein interaction networks [19]. There are several reasons why: First, because they encode proteins required for universal and continuous cellular processes such as transcription, translation, and energy metabolism, they function within complex molecular pathways which would require numerous physical interactions [20]. Their broad expression across all cell types also increases both true interaction opportunities and the observed connectivity due to higher cellular abundance. In other words, since these genes are found everywhere, they have more places to make interactions and are more likely to be well-studied. Finally, HK genes are typically evolutionarily ancient and conserved, thus they have had more time to accumulate interactions with other proteins [20,21]. These reasons likely underlie our findings that housekeeping genes were disproportionately allocated to the Support Group compared with other model-selected groups at both temperatures. Note that the Support Group was created based on analyzing inter-individual variability, completely independent from degree, but the results indicate these genes are more well-connected (have a higher median degree) than the background and other DEGs at both temperatures. After excluding known housekeeping genes, neither Support Group degree comparison remained significant, suggesting that housekeeping members underlie the elevated degree pattern. Support genes are the set of genes that have high inter-individual variability both before and after treatment (definition 2.1.3 in [4]). Gonzalez *et al.* (2026) assert that these are still well-regulated genes in the sense that their behaviors are consistent before and after treatment [4]. It was assumed that their variability remained consistently large because they are connected to many biological processes. Our results substantiate this assumption. They indicate that we may be able to define high degree/housekeeping genes in terms of connectivity and in terms of variability since they can have a similar likelihood of being found in the known HK database.

Wolf et al. (2023) examined inter-individual expression variation across 57 human RNA-seq datasets from nonperturbed conditions [22]. They found that low-variance genes were enriched for housekeeping function and that expression variance was weakly negatively associated with connectivity, measured through co-expression correlations. In our study, the Support Group (g3) had both higher connectivity, measured by documented protein interactions, and enriched housekeeping overlap. Both studies associate housekeeping function with higher connectivity, despite measuring connectivity differently. There is a contrast, however, regarding expression variability: housekeeping genes were enriched among Wolf et al.’s low-variance genes, whereas our greater housekeeping overlap occurred in a group defined by high variability both before and after temperature perturbations. While Wolf and colleagues analyzed variation across individuals under nonperturbed conditions in multiple tissues, we examined fibroblasts before and after temperature treatment. The generalizability of the overlap we observed between housekeeping genes and high expression variability before and after environmental perturbations therefore remains unclear.

Additionally, while we compared our network-building algorithm to STRING-db’s, they do rely on slightly different proteomic data. Our network-builder relied on Human Protein Atlas’s compilation of literature-curated physical interactions whereas STRING includes broader functional associations such as co-expression, text mining, neighborhood, gene fusion, and co-occurrence.

An inherent limitation to using both STRING-db and our framework to test gene sets is that protein interaction data capture only part of the cell’s response to an environmental perturbation. Future work could include pathway/reaction data involving metabolites and lipids from Reactome or KEGG and RNA-associated interactions from RNAInter to make more granular predictions of which differentially expressed genes warrant further inquiry [23–25]. However, incorporating these relationships would complicate the DEG-filter of our networks since metabolites and lipids are not directly encoded by individual genes. Moreover, the goal of this paper is not to model the cell’s comprehensive mechanistic response to perturbations, but rather to develop a framework for prioritizing specific DEG subsets based on proteomic data.

## Conclusion

We provide a novel algorithm that uses database-curated protein-protein interactions to evaluate differentially expressed gene sets from RNA-seq data. By restricting shared second-order interaction intermediaries to DEGs, we aim to add a condition-informed filter to protein-protein interaction networks. Applied to temperature-response gene sets from Gonzalez *et al.* (2026) and *limma*, the algorithm identified larger-than-expected DEG-restricted LCCs at 32°C [4]. The 32°C Principal Response genes also retained significantly more LCC members under DEG restriction than expected from random sets, indicating that this DEG-selection model appears to choose genes that are biologically meaningful and assemble into coherent networks, a prerequisite to enacting a coordinated response to environmental change [4]. Corresponding DEG-restricted LCC-size and connectivity-retention comparisons at 41°C were not significant.

Beyond the Principal Response findings, we also found that housekeeping genes were disproportionately represented in G3 compared with the other model-selected groups at both temperatures, with significant enrichment relative to the broader background at 41°C.

Attempting to increase specificity by restricting second-order interaction intermediates to proposed DEGs may offer more precise and context-informed validation of future studies. As noted, our algorithm should only serve as a preliminary tool before further gold-standard experimental analyses are undertaken to test how model-identified genes act together in response to environmental perturbation or differential phenotypes. Future work would build on these results by validating more detailed individual model predictions against the biological literature and distinguishing between positive and negative forms of protein-protein interactions. Expanding the framework to other computational models of gene regulation under different types of perturbations will further test its generalizability. In the long term, this work contributes to establishing more reliable approaches for integrating computational modeling with experimental biology, which has implications for stress physiology and computational modeling of genomic responses to environmental changes.

## DATA AVAILABILITY

All source code and input files are available at https://github.com/BradleyFrishman/Context-Specific-DEG-PPI-Network-Builder. Public databases used for this study can be accessed at https://www.proteinatlas.org/ and https://string-db.org/.

## ACKNOWLEDGMENTS

The authors would like to thank Diane Genereux and the NSF Flexible Homeostasis team for the opportunity to present this project at their annual group meeting and for the helpful feedback. The authors acknowledge the use of generative artificial intelligence (AI) tools, including ChatGPT (OpenAI), in the development of the Python scripts used in this study. All final code design, methodological decisions, biological interpretation, and validation of outputs were performed by the authors. A preprint is available at https://doi.org/10.64898/2025.12.10.693541

## AUTHOR CONTRIBUTIONS

Bradley Frishman (Conceptualization, Formal analysis, Methodology, Writing original draft), Jorge Gonzalez (Conceptualization, Formal analysis, Methodology, Writing – review and editing), Valery Forbes (Conceptualization, Supervision, Writing – review and editing). All authors read and approved the final manuscript.

