## Supplementary Information for "A Condition-Informed Framework for Evaluating Temperature-Responsive Gene Sets Using Protein Interaction Data"

### Degree-Shifted Reference Scenarios/Effect Size Analysis

To gain a better understanding of how significant a small observed difference in the LCC size may be, we performed degree-shifted reference scenarios. For each temperature condition, we first computed the degree distributions of the Principal Response genes, defined as the number of known interactions per gene. We then generated four modified degree distributions. In the conservative upshift, only the lowest non-zero-degree bin was moved up by one. That is, we found the lowest degree bin and moved all the genes counted in that bin to the next higher bin. In the aggressive upshift, the entire degree distribution was shifted one bin to the right. The same logic applies for the converse. In the conservative *downshift*, we again moved only the (number of) genes in the lowest bin down into an even lower degree bin. In the aggressive downshift, we shifted the entire distribution down by one bin. For each scenario, we repeatedly sampled background genes (this refers to the 9825 candidate genes) to match the target degree distribution as closely as possible, built PPI networks, and measured LCC.

After performing degree-shifted reference scenarios, we found that the genes selected to be used in the model (for either temperature condition) tended to have a larger LCC even when compared to a gene set with conservatively more known interactions (**Figure S1**). The purpose of this analysis was to bound the LCC and to gain insight into how sensitive it is to conservative or aggressive changes in degree distribution. Using this information, we can better assess how significant the apparent differences between LCCs are.

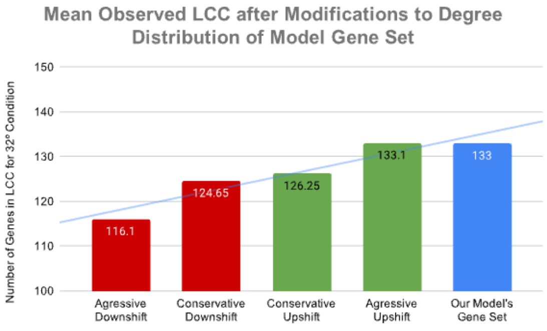


(A)

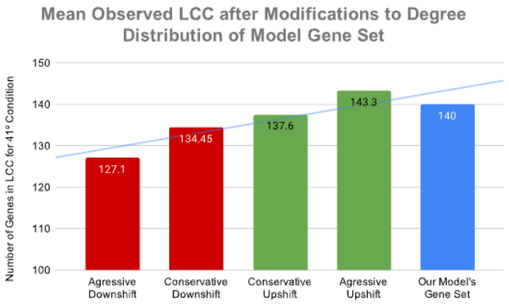

(B)

Supplementary Figure S1. LCC Change After Distribution Shifts. (A) represents the results from the 32º treatment, (B) represents the results from the 41º treatment. 20 iterations per scenario without a random seed were used for this analysis.

### Further Support Group (G3) Genes Analysis

| G3 set | Genes | G3 median degree | Background median | Unadjusted Empirical p |
|---|---|---|---|---|
| 32°C all | 67 | 22 | 13 | **0.016** |
| 32°C non-HK | 49 | 16 | 10 | 0.073 |
| 41°C all | 53 | 22 | 13 | **0.026** |
| 41°C non-HK | 35 | 9 | 10 | 0.72 |

Supplementary Table S1. Median degree was compared with 1,000 size-matched random sets. All-gene tests use the 9,825-gene background; non-HK tests exclude HK atlas members from both G3 and the background (7,686 genes). These tests intentionally do not degree-match the random sets. **Bold indicates p < 0.05.**

| Group | 32° | 41° |
|---|---|---|
| G3 Housekeeping Percentage [HK / (HK + non-HK)] | (18 / 67) = 26.87% | (18 / 53) = 33.96% |
| Other model groups G1+2+4 Housekeeping Percentage [HK / (HK + non-HK)] | (20 / 182) = 10.99% | (24 / 204) = 11.76% |
| G3 vs. other model groups: Fisher p | **0.0027** | **0.00027** |
| Background outside G3 Housekeeping Percentage [HK / (HK + non-HK)] | (2,121 / 9,758) = 21.74% | (2,121 / 9,772) = 21.70% |
| G3 vs. background outside G3: Fisher p | 0.19 | **0.028** |

Supplementary Table S2. Fisher contingency counts. At both temperatures, housekeeping genes are more concentrated in G3 than in the other selected groups. Against the broader background, G3 contains significantly more housekeeping genes than expected from the eligible gene pool at 41°C, but not at 32°C. **Bold indicates $p < 0.05$.**

(a)

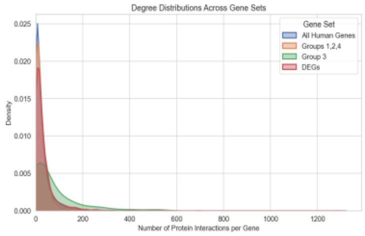


(b)

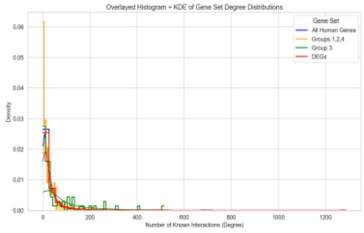


(c)

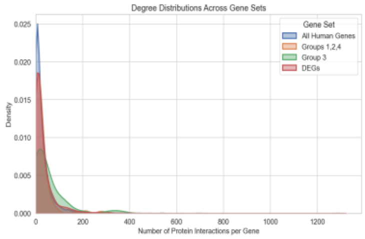


(d)

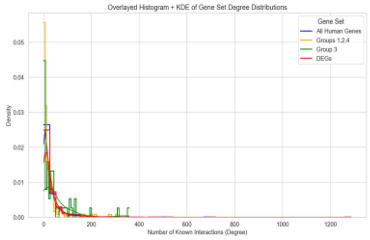


Supplementary Figure S2. Analysis of Gonzalez Model Selected Group 3 Genes' Number of Known Interactions (Degree). These statistics indicate that the model-categorized Group 3 (Support Group) Genes have relatively more known interactions compared to all candidate genes, proposed DEGs, and the other groups of proposed DEGs.

a. 32º Kernel Density Estimation (KDE) with the x-axis representing the number of known interactors for each gene, based on Human Protein Atlas data. The Y-axis ("Density") reflects the relative frequency (via a kernel density estimate) of genes with a given number of interactions within each group. Normalized so the area under each curve equals 1 (not absolute counts).
b. 32º Histograms and KDE.
c. 41º. KDE with the x-axis representing the number of known interactors for each gene, based on Protein Atlas data. The Y-axis ("Density") reflects the relative frequency (via a kernel density estimate) of genes with a given number of interactions within each group. Normalized so the area under each curve equals 1 (not absolute counts).
d. 41º Histograms and KDE.

| | All 9825 Human Genes Studied | 32º DEGs | 32º Groups 1,2,4 | 32º Support Group/ Group 3 (Suspected Housekeeping) | 41º DEGs | 41º Groups 1,2,4 | 41º Support Group/ Group 3 (Suspected Housekeeping) |
|---|---|---|---|---|---|---|---|
| Median Degree | 13.0 | 15.0 | 12.0 | 22.0 | 16.0 | 11.0 | 22.0 |
| Mean Degree | 28.12 | 31.77 | 22.47 | 66.22 | 32.67 | 24.34 | 48.68 |
| Std Dev Degree | 48.98 | 55.84 | 29.97 | 100.65 | 56.22 | 44.77 | 72.21 |

Supplementary Table S3. Statistical Analysis of the Varying Degree Distributions. These statistics quantitatively show that the model-categorized Group 3 (Support Group) Genes have relatively more known interactions compared to all candidate genes, proposed DEGs, and the other groups of proposed DEGs.

#### Second order LCC size comparison

| Gene set | Unrestricted LCC | Restricted LCC | Unrestricted p | Restricted p |
|---|---|---|---|---|
| Principal Response 32°C | 153 | 133 | 0.7033 | **0.0250** |
| Principal Response 41°C | 175 | 140 | **0.0490** | 0.1229 |
| G3 32°C | 61 | 57 | **0.0100** | **0.0030** |
| G3 41°C | 44 | 36 | 0.1219 | 0.4316 |
| limma Principal-Response-sized subset 32°C | 169 | 149 | **0.0010** | **0.0070** |
| limma Principal-Response-sized subset 41°C | 184 | 142 | **0.0020** | 0.6234 |
| limma G3-sized subset 32°C | 58 | 42 | 0.2048 | 0.6723 |
| limma G3-sized subset 41°C | 45 | 25 | 0.0809 | 0.8462 |

Supplementary Table S4. Restricted and unrestricted second-order networks use the same cached interactions and paired random gene sets. Each p compares the observed LCC with its own random-network distribution. P-values unadjusted. **Bold indicates $p < 0.05$.**

### Corresponding Limma Subset of Genes to Gonzalez's G3/Support Group

| Condition | Initially # genes selected | Retained # genes for network analyses below |
|---|---|---|
| 32°C | 67 (to match the number of Group 3 genes) | 63 |
| 41°C | 53 (to match the number of Group 3 genes) | 51 |

Supplementary Table S5. Initial number of genes selected to match the number of genes in G3/Support Group and the retained number of genes for network analyses below. 4 genes in the initial 32ºC set had no known interactions in our interaction cache from Proteinatlas.org and 2 genes in the initial 41º set had no known interactions in our interaction cache from Proteinatlas.org, so they were excluded from the analyses seen below.

DEG-restricted networks for Corresponding Limma Subset of Genes to Gonzalez's G3/Support Group:

| Condition | Network | LCC size | Null mean size | Size p | LCC edges | Null mean edges | Edge p |
|---|---|---|---|---|---|---|---|
| 32°C | Second-order only | 42 | 43.0 | 0.6723 | 152 | 133.8 | 0.2607 |
| 32°C | First + second order | 43 | 43.6 | 0.6374 | 154 | 135.7 | 0.2597 |
| 41°C | Second-order only | 25 | 28.0 | 0.8462 | 55 | 64.3 | 0.6863 |
| 41°C | First + second order | 25 | 28.5 | 0.8731 | 60 | 65.3 | 0.5914 |

Supplementary Table S6. LCC size and LCC edge count were not significantly larger or smaller than expected for the corresponding Limma subset of genes to G3/Support Group genes. P-values unadjusted.

### Connectivity retention after DEG restriction for Corresponding Limma Subset of Genes to Gonzalez's G3/Support Group:

| Condition | Network | Unrestricted LCC | Restricted LCC | Observed loss (ΔLCC) | Mean random loss (ΔLCC) | Smaller-loss p |
|---|---|---|---|---|---|---|
| 32°C | Second-order only | 58 | 42 | 16 | 13.0 | 0.8322 |
| 32°C | First + second order | 58 | 43 | 15 | 12.5 | 0.7952 |
| 41°C | Second-order only | 45 | 25 | 20 | 13.6 | 0.9500 |

| | | | | | | |
|---|---|---|---|---|---|---|
| 41°C | First + second order | 45 | 25 | 20 | 13.2 | 0.9610 |

Supplementary Table S7. LCC member retention was not significantly larger or smaller than expected after applying our DEG filter to second-order intermediaries for the corresponding Limma subset of genes to G3/Support Group. P-values unadjusted.

### Unrestricted networks for Corresponding Limma Subset of Genes to Gonzalez's G3/Support Group:

| Condition | Network | LCC size | Null mean LCC size | LCC Size p | LCC edges | Null mean LCC edges | LCC Edge p |
|---|---|---|---|---|---|---|---|
| 32°C | Second-order only | 58 | 56.0 | 0.2048 | 384 | 298.6 | **0.0180** |
| 32°C | First + second order | 58 | 56.1 | 0.2148 | 384 | 299.5 | **0.0190** |
| 41°C | Second-order only | 45 | 41.6 | 0.0809 | 138 | 154.0 | 0.7303 |
| 41°C | First + second order | 45 | 41.7 | 0.0839 | 140 | 154.5 | 0.7103 |

Supplementary Table S8. Unadjusted enrichment p-values are upper-tail empirical tests. **Bold indicates p < 0.05**. Only the 32ºC corresponding limma subset of genes to Gonzalez's G3/Support Group edge count showed significance within their LCC before applying our DEG-filter to second-order intermediaries in the PPI network.

### **Sensitivity of empirical network-enrichment p-values to the number of randomizations.**

Significance classifications were unchanged across 500, 1,000, and 1,500 randomizations for all LCC-size and edge-count comparisons. Two comparisons changed classification between 100 and 500 randomizations: combined-network LCC size for the 32°C Principal Response and limma sets. For both 32°C G3 network definitions, no random LCC matched or exceeded the observed edge count at any checkpoint; decreasing p-values therefore reflected the finer resolution available with additional randomizations (Supplementary Table S4).

| **Gene set** | **Network** | **Statistic** | **100** Randomizations p | **500** Randomizations p | **1,000** Randomizations p | **1,500** Randomizations p |
|---|---|---|---|---|---|---|
| Principal Response 32°C | Second-order only | LCC size | **0.0495** | **0.0259** | **0.0250** | **0.0280** |
| Principal Response 32°C | First + second order | LCC size | 0.0693 | **0.0479** | **0.0480** | **0.0493** |
| Principal Response 41°C | Second-order only | LCC size | 0.1287 | 0.1257 | 0.1229 | 0.1292 |
| Principal Response 41°C | First + second order | LCC size | 0.2178 | 0.1956 | 0.1918 | 0.2005 |
| G3 32°C | Second-order only | LCC size | **0.0099** | **0.0040** | **0.0030** | **0.0027** |
| G3 32°C | First + second order | LCC size | **0.0099** | **0.0060** | **0.0040** | **0.0053** |
| G3 41°C | Second-order only | LCC size | 0.3861 | 0.4331 | 0.4316 | 0.4377 |
| G3 41°C | First + second order | LCC size | 0.3267 | 0.3413 | 0.3257 | 0.3431 |
| limma 32°C | Second-order only | LCC size | **0.0099** | **0.0040** | **0.0070** | **0.0080** |
| limma 32°C | First + second order | LCC size | 0.0594 | **0.0299** | **0.0280** | **0.0266** |
| limma 41°C | Second-order only | LCC size | 0.6832 | 0.6367 | 0.6234 | 0.6302 |
| limma 41°C | First + second order | LCC size | 0.8020 | 0.8004 | 0.7872 | 0.7921 |

| | | | | | | |
|---|---|---|---|---|---|---|
| G3 32°C | Second-order only | LCC edge count | **0.0099** | **0.0020** | **0.0010** | **0.0007** |
| G3 32°C | First + second order | LCC edge count | **0.0099** | **0.0020** | **0.0010** | **0.0007** |
| G3 41°C | Second-order only | LCC edge count | 0.2277 | 0.1956 | 0.1998 | 0.1979 |
| G3 41°C | First + second order | LCC edge count | 0.1980 | 0.1856 | 0.1888 | 0.1865 |

Supplementary Table S9. Upper-tail unadjusted empirical p-values for DEG-restricted networks were calculated as p = (b+1)/(N+1), where b is the number of random networks with a statistic greater than or equal to the observed value and $N$ is the number of randomizations. Results use the first 100, 500, 1,000, and 1,500 draws from the same randomization sequence for each gene set, rather than independent replicate runs. Random sets were size-matched and approximately degree-matched. **Bold indicates p < 0.05.**
